# Selection-based model of prokaryote pangenomes

**DOI:** 10.1101/782573

**Authors:** Maria Rosa Domingo-Sananes, James O. McInerney

## Abstract

The genomes of different individuals of the same prokaryote species can vary widely in gene content, displaying different proportions of core genes, which are present in all genomes, and accessory genes, whose presence varies between genomes. Together, these core and accessory genes make up a species’ pangenome. The reasons behind this extensive diversity in gene content remain elusive, and there is an ongoing debate about the contribution of accessory genes to fitness, that is, whether their presence is on average advantageous, neutral, or deleterious. In order to explore this issue, we developed a mathematical model to simulate the gene content of prokaryote genomes and pangenomes. Our model focuses on testing how the fitness effects of genes and their rates of gene gain and loss would affect the properties of pangenomes. We first show that pangenomes with large numbers of low-frequency genes can arise due to the gain and loss of neutral and nearly neutral genes in a population. However, pangenomes with large numbers of highly beneficial, low-frequency genes can arise as a consequence of genotype-by-environment interactions when multiple niches are available to a species. Finally, pangenomes can arise, irrespective of the fitness effect of the gained and lost genes, as long as gene gain and loss rates are high. We argue that in order to understand the contribution of different mechanisms to pangenome diversity, it is crucial to have empirical information on population structure, gene-by-environment interactions, the distributions of fitness effects and rates of gene gain and loss in different prokaryote groups.

## Introduction

Intraspecific variation in gene content in prokaryotes is often so extensive that for a given species a “typical genome” might not exist^1,2^. Therefore, the concept of a pangenome, defined as the complete set of genes present in a species, has been put forward as a better description of the gene content of a prokaryote species^1^. Pangenomes include *core* genes that are present in all individuals, and *accessory* genes, whose presence varies among individuals. The role of accessory genes remains puzzling, and a debate has developed centring on whether they have positive, negative, or neutral effects on the cells that host them^3–8^. Understanding the contributions of these accessory genes to cell fitness and evolution has important implications, including insights into the factors that determine the prevalence and maintenance of virulence genes, antibiotic resistance, and other traits often encoded by accessory genes^1,9^.

Pangenomes arise as a consequence of gene acquisition via horizontal gene transfer (HGT), and gene loss^2^. These processes ultimately result in general patterns observed across prokaryotes, which include an increase in the number of known accessory genes as more genomes from the same species are sequenced —along with a slower decrease in the number of core genes— and a U-shaped *gene frequency distribution* or *spectrum*^2,10,11^. The U shape of this distribution indicates that there is a large proportion of genes present in a single or very few genomes, few genes present at intermediate frequencies, and substantial proportion of core genes. Although across prokaryotic life there is a certain amount of commonality in the shape of the gene content frequency distributions, different prokaryote species manifest significant differences in the proportions of core and accessory genes^12–14^.

Although we know that HGT and gene loss result in pangenomes, what is less clear are the forces that lead to high diversity in gene content and U-shaped gene frequency distributions. According to mathematical models, gain and loss of entirely neutral genes along a simulated phylogeny or population can in principle predict the U-shaped gene frequency distributions of pangenomes ^11,15^. However, the fit to real data improves significantly when at least two classes of genes, with either slow or fast turnover rates, are considered^11,15,16^. Specifically, including a class of genes with low turnover rates increases the number of core and very common genes, thus allowing a better fit to the peak of high frequency genes in the distribution. This implies that selection is required to retain core genes for long periods of time, exactly as we would expect if these have important functions. Therefore, these models suggest that high frequency genes are beneficial on average, while most accessory and very rare genes are likely to be neutral.

Alternative models suggest that accessory genes are on average beneficial, and propose selection-based mechanisms that can explain the maintenance of diversity in gene content^12,17–19^. One possibility is that negative-frequency dependent selection (NFDS) acting on large numbers of accessory genes explains the existence and maintenance of gene content variation^20,21^. NFDS occurs when the fitness effect of a gene changes from positive to negative as its frequency in the population increases. That is, the gene is beneficial when rare but deleterious when common and is therefore maintained at an intermediate frequency in the population. NFDS may be important for maintaining gene content diversity under some conditions, such as when rare serotypes become fitter after a vaccination program that only targets common serotypes^17,18^. A related possibility is the existence of Black Queen effects, where the combination of pressure to reduce genome size while maintaining common-good functions results in intermediate frequencies of the genes encoding such functions^22^. Genotype-by-environment (GxE) effects have also been proposed to maintain beneficial genes at an intermediate frequency in a population^12^. GxE effects imply that particular genes are beneficial only in some conditions and are therefore only maintained in particular environments. Migration of cells between different ecological niches, or distinct environments, could explain variation in gene content and the maintenance of genes at low frequencies due to differences in their fitness in the different environments^19^.

Overall, different models propose that the existence of pangenomes can be the consequence of either neutral processes or selection. We do not currently know which of these models better explains pangenomes in nature. Most likely, genes with deleterious, neutral and beneficial effects are all present in the accessory component of pangenomes. The more difficult question is whether we can determine which fitness class is the most prevalent and, at the same time, understand what processes and pressures determine the existence and maintenance of pangenomes and their properties. Here we develop a mathematical model from the point of view of a single gene that has a particular fitness contribution (that can be positive or negative) and that can be gained or lost. We analyse the consequences that variation in fitness effects, and the rates of gene gain and loss, have on the expected frequencies of a gene in a population where drift is negligible. We then expand the single-gene model in order to consider many genes, each allowed to have different properties and using this genome-scale expansion, we simulate pangenomes and their gene frequency distributions. We show that the distributions of the input parameters directly influence pangenome properties such as the size of the accessory genome, core genome and the shape of the gene frequency distribution. We also model the effect that multiple niches, and niche-dependent gene fitness contributions have on pangenomes. Our results show that the presence of accessory genes is likely to be overwhelmingly nearly neutral in the absence of GxE effects, which is likely the case when populations are constrained to a single environmental niche (*i.e.* there is no migration or exposure to variation in environmental conditions). However, when multiple environmental niches are considered, accessory genes can be highly beneficial. Overall, we argue that real pangenomes are an outcome of a combination of processes and crucially, variability in model parameters, like fitness effects and rates of gene gain and loss, for different genes and species.

## Model description

We propose the model depicted in Figure 1a for the frequency of a gene that can be present or absent in the genomes of a prokaryote group (for example a population or species). We assume that cells without the gene divide at rate *μ*, and that the gene’s presence provides a contribution *s* to this growth rate, such that cells that possess the gene divide at rate *μ* + *s*. We call *s* the selection coefficient or *fitness effect* of the gene, and its value can be positive or negative. For cells that lack the gene, it can be gained at a rate *r*_*g*_, and lost from cells that have it with rate *r*_*l*_ (Figure 1a). We assume that the population size is constant and that all cells in the population have the same death rate *v* for cells with and without the gene. We also assume that genetic drift in the population is negligible and that the only changes in genomes are the gain and loss of genes. That is, there are no other mutations (that would, for example, change the value of *s*). We can then write a differential equation for the frequency of the gene in the population, *x*:

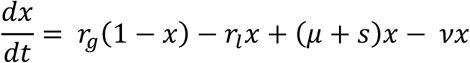

**Figure 1:**
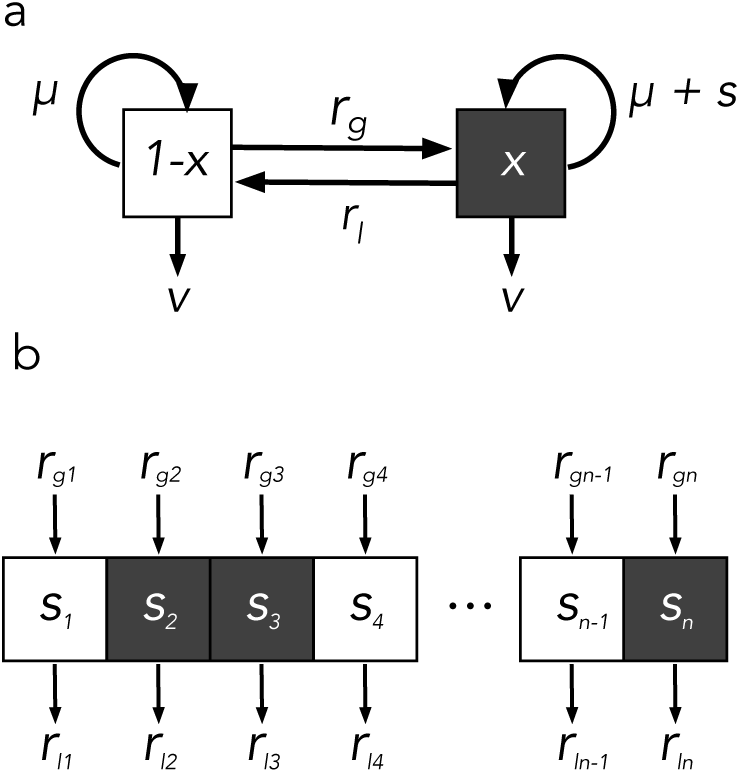
Selection-based model of pangenomes. (a) Model scheme for the frequency of one gene. (b) Model scheme for multiple genes and pangenome simulations.

where

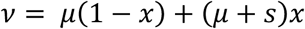

because the population size is constant (*d*(*x* + (1 − *x*))/*dt* = 0). Therefore, we obtain the following equation, which describes the frequency of the gene independently of the basal growth rate of the population:

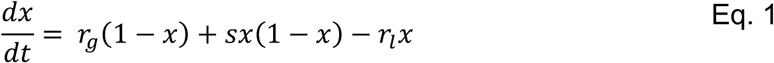

Thus, in this model, the frequency of a gene in a population depends on three parameters: its fitness effect (*s*), its rate of gain (*r*_*g*_) and its rate of loss (*r*_*l*_). These parameters summarise important features of the external environment, the recipient cell, and the gene itself:

The *fitness effect* of a gene describes the additive contribution of that gene to fitness, or more narrowly to the cell’s growth rate. That is, it describes the difference in growth provided by the gene per unit time. Therefore, *s* > 0 if the gene confers a trait that increases fitness, (i.e. if it is beneficial), *s* < 0 if the gene’s presence is deleterious, and *s* = 0 if the gene has a neutral effect with respect to fitness. The fitness effect is not just a property of the gene itself but depends on the organism’s biology. For example, it depends on whether or not the cell can make use of that gene, in terms of compatibility with its cell biology and codon usage, for instance. Importantly, the value of this parameter also depends on the environment. For example, a gene that allows a cell to metabolise a substrate, or protect it from an antibiotic, will only be beneficial in the relevant external ecological setting.

The *rate of gain* for a given gene describes the proportion of cells in the population that gain that gene per unit of time. The value of this parameter is heavily dependent on the route through which the gene can be acquired. For example, genes acquired by transduction or conjugation are likely to have high rates of gain, while those acquired through natural transformation will have lower gain rates. Importantly, this parameter also depends on the prevalence of a gene in the environment (whether it is environmental DNA, phage, plasmids or other cells), because the higher the prevalence, the higher the chance a gene can be acquired by a cell in the focal population. Finally, the rate of gain is also dependent on the characteristics of the species or cells where the gene is acquired. For example, factors such as the intrinsic rates of natural transformation and recombination, the prevalence of lysogenic phage infections and conjugative plasmids, and even the existence and efficiency of defence mechanisms that can prevent the entry of exogenous DNA, will impact on the gain rate of genes. The *rate of loss* for a given gene is the proportion of cells that lose that gene per unit time, and likely depends mostly on the organism’s biology, i.e. rates of recombination that lead to gene loss. However, this rate may also vary with the environment and properties that enhance or reduce rates or recombination of particular genes, such as the types of sequences flanking the gene^23,24^.

Our model does not explicitly include evolutionary history (*i.e.* phylogeny) and drift, therefore, the implication is that selection is quite effective. Consequently, our model is at the opposite end of the spectrum of assumptions when compared to models based exclusively on neutral evolution^11,15^. We use this model to explore how the values of these parameters and their distributions determine genome content and pangenome properties.

## Results

### Effect of model parameters on pangenome gene frequencies

According to the model (Eq 2, Methods) the steady state gene frequency in a prokaryote population is a sigmoid function of the fitness effect *s*, which means that the frequency of a gene increases monotonically from 0 to 1 as the value of *s* increases, but changes in a switch-like manner around *s* = 0 (Figure 2a). Overall, this implies that deleterious genes are present at low frequencies, while beneficial genes are found at high frequencies. The rates of gene gain and loss influence the shape of this sigmoid function. As the gain and loss rates decrease, the sharpness or switch-like shape of the function increases (compare purple and blue lines in Figure 2a). The consequence of lower rates of gene gain and loss is a decrease in the frequency of deleterious genes, and an increase in the frequency of beneficial genes. In the limit where the gene gain and loss rates are equal to zero, we obtain a perfect step function (Equation 2), such that all advantageous genes are fixed in the population and all deleterious genes are absent from the population, even if their fitness effects are very small (due to our assumption of strong selection). This result implies that at equilibrium, fitness is maximised when the rates of gene gain and loss are very small compared to fitness effects. However, if the gene gain and loss rates are equal to zero, any beneficial genes not already present in the genome cannot be gained, while any deleterious genes cannot be lost. Additionally, if the rates of gene gain and loss are very small, it takes longer to reach steady state frequencies and the gene frequency dynamics are mainly driven by selection (see time-course simulations in Figure S1). Furthermore, in agreement with evolutionary theory, the smaller the fitness effect, the slower the change in gene frequency (not shown).

**Figure 2.**
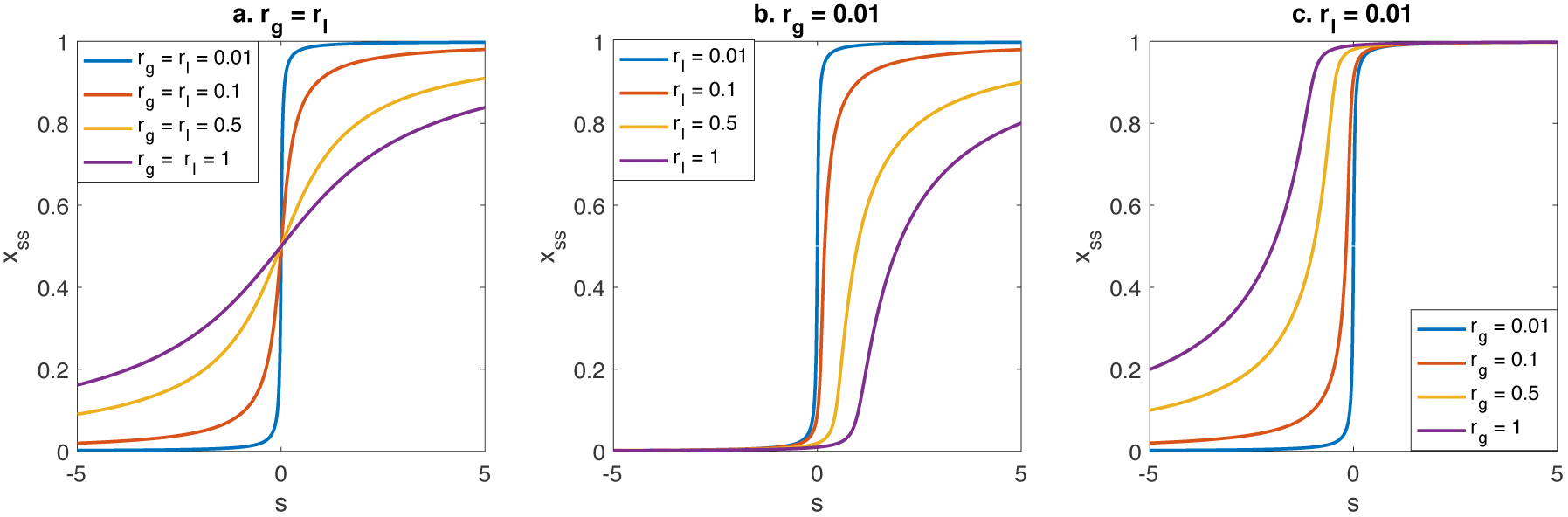
Dependence of gene frequency with respect to model parameters. Plots shows the gene frequency with respect to the fitness effect, *s*, for different rates of gene gain and loss. A. Effect of rates of gene and loss when *r*_*g*_ = *r*_*l*_. B. Effect of the rate of gene loss with fixed rate of gene gain (*r*_*g*_ = 0.01). C. Effect of the rate of gene gain with fixed rate of gene loss (*r*_*l*_ = 0.01).

The relative balance between the rates of gene gain and loss is also important for determining the equilibrium gene frequency. Figure 2b shows that if the rate of gene loss increases (from blue to purple line) while the rate of gain remains fixed, the frequency of deleterious genes remains low, while the frequency of beneficial and nearly neutral genes decreases. Therefore, according to the model, if rates of loss are higher than rates of gain, deleterious genes are efficiently purged from the population. The drawback to higher rates of loss is that highly beneficial genes can also be lost at high rates and are therefore not fixed in the population at equilibrium. The disparity in the rates of gain and loss also has an effect on gene frequency dynamics. At high loss rates, deleterious genes are quickly lost, but acquisition of advantageous genes is delayed in comparison with the case of equal gain and loss rates (Figure S1). Essentially the opposite situation occurs if the rates of gain are higher than the rates of loss: beneficial genes will be fixed in the population, but neutral and deleterious genes will also be present at high frequencies (Figure 2c). Furthermore, the model shows that highly deleterious genes will be present at low or intermediate frequencies. In terms of dynamics, higher rates of gene gain than loss also implies that advantageous genes quickly fix in the population, but deleterious genes are lost slowly (Figure S1).

Overall, according to our model, it is not just the fitness effects of accessory genes that can determine the frequency of a gene in the population, but the rates of gene gain and loss are also important, especially if their values are of a magnitude comparable to the fitness effects of genes. In particular, the frequency of nearly neutral genes in a population is heavily dependent on the balance between rates: if gain rates are higher than loss rates, neutral genes are maintained in the population at high frequency, while they will be rare when loss rates are higher (Figure S2).

### Effect of parameter distributions on pangenomes

In a population of cells in which multiple genes can be gained and lost, the values of the model parameters are likely to vary among different genes. Furthermore, even for a single gene, its fitness effect, rate of gain and rate of loss can vary among different species, cells and environments. Therefore, each of the gene-specific model parameters is associated with a parameter distribution. Some point estimates for rates of gene gain and loss have been obtained^25^ and individual examples of fitness effects of accessory genes are known^8,26^ however, there is very little empirical data on these parameter distributions. Consequently, in our model we rely on hypothetical parameter distributions in order to assess how parameter variation shapes pangenomes. We sampled gene parameter values from proposed distributions and simulated gene content for multiple genomes in a population at equilibrium as described in Methods. Importantly, because we know the parameter values of these simulated pangenomes, we assessed the contribution to pangenomes of genes with different fitness effects.

#### Distribution of fitness effects

The distribution of fitness effects (DFE) of genes is central to our expectations of the gene content variation we see in populations. This distribution is analogous to the distribution of fitness effects of mutations, which is important in determining the level of nucleotide polymorphisms that are maintained in populations^8,27^. We currently know little about the DFE of accessory genes, but as a first approximation we speculate that it may have similar properties to the DFE for mutations: most genes will have nearly neutral effects and highly beneficial or highly deleterious genes will be rare.

As a first approximation, we assume equal and relatively low rates of gene gain and loss for all genes. For simplicity, we also assume a finite pool of genes available for acquisition. Figure 3A shows results from a simulation of 100 genomes, sampled from a gene pool of 1000 genes whose fitness effects are drawn from a normal distribution with a mean of zero and unit standard deviation (blue density plot in Figure 3Ib). The top panel (Figure 3Ia) shows the presence/absence of all the genes in the 100 simulated strains as a heatmap, and the scatter plot (Figure 3Id) shows the frequency of every gene in the pangenome with respect to its fitness effect. In this case, advantageous genes (*s* > 0) are present at high frequencies, nearly neutral genes present at intermediate frequencies and deleterious genes (*s* < 0) are very rare or absent. This simulation thus qualitatively recovers a U-shaped gene frequency distribution similar to those observed for real pangenomes^2^ (Figure 3Ic). In these simulations highly advantageous genes are essentially fixed and would be considered the ‘core’ genome, while nearly neutral genes and some deleterious genes would constitute the ‘accessory’ genome. This also results in ‘filtering’ of a fraction of the deleterious genes, such that the fitness effects of genes present in the pangenome (orange density plot in Figure 3Ib) are slightly higher than the original gene pool DFE. These results demonstrate that the pattern of gene content variation of real pangenomes can be recovered by a model based on selection only. This contrasts with neutral models, which propose that the U-shaped gene frequency distributions observed in prokaryote pangenomes can be explained by entirely neutral genes subject only to drift in an evolving population^11,15^. Therefore, models with opposite assumptions regarding the effects of drift and selection, are able to qualitatively recover patterns of gene content variation that are similar to empirically observed patterns.

**Figure 3.**
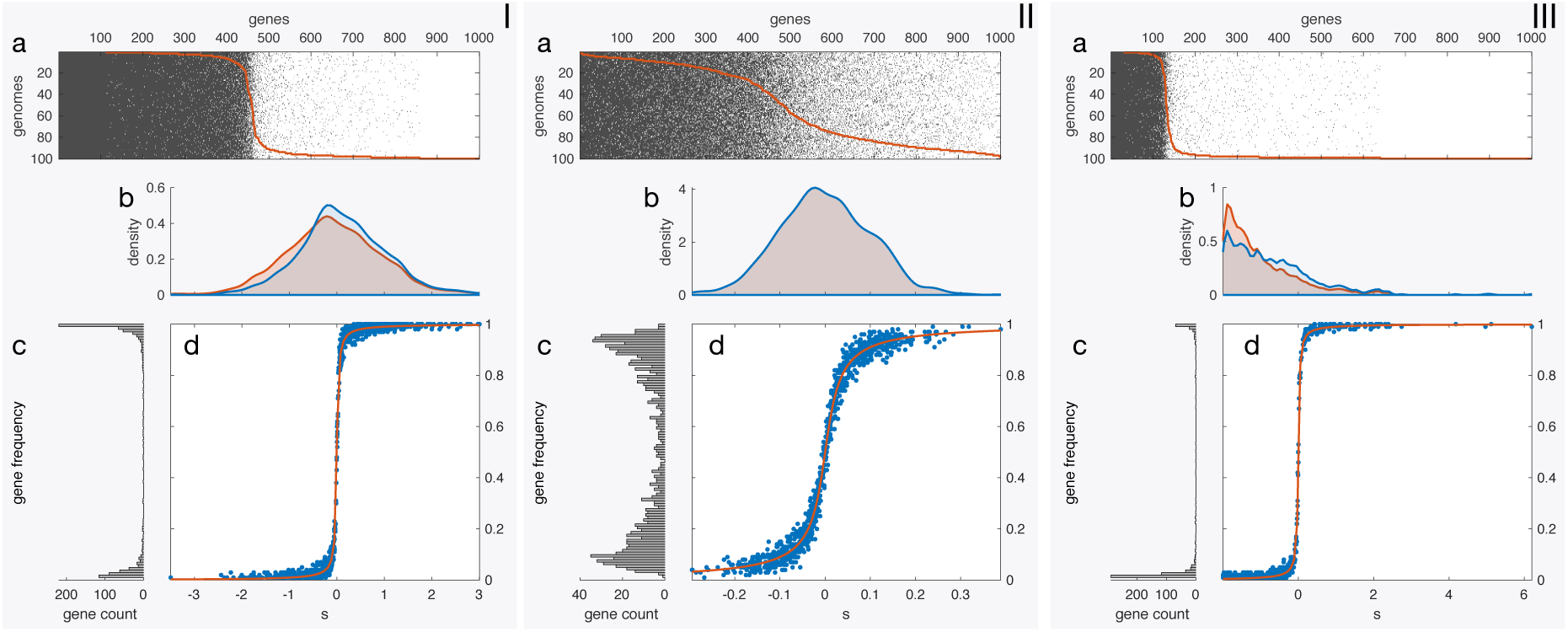
Simulated pangenomes for different distributions of fitness effects. Each panel represents a simulation with fitness effects sampled from the specified distribution of fitness effects. **I**. Normal distribution with mean 0 and SD = 1. **II**. Normal distribution with mean 0 and SD = 0.1. **III**. Exponential distribution with rate 1, mean shifted to −1. For each panel: **a**. Presence/absence of genes in the simulated pangenome, present genes are indicated in grey. The orange line indicates the number of strains in which a gene is present. **b**. Density plots for the DFE of the gene pool (sampled from the specified theoretical distribution, in blue) and for the fitness effects of genes actually present in the pangenome (orange). **c**. Gene frequency distribution for the simulated pangenome. **d**. Gene frequencies versus fitness effects for 1000 genes in the simulated pangenome (blue dots) and the theoretical relationship (orange line). Rates of gene gain and loss are assumed to be equal for all genes (*r*_*g*_ = *r*_*l*_ = 0.01)

In our model, if we assume most genes are nearly neutral, the gene frequency distribution starts to deviate from the expected U shape. Compared to the model with larger variance in fitness effects (Figure 3I), if we significantly reduce the magnitude of fitness effects of the DFE (∼10-fold reduction) the result is that more genes are present at intermediate frequencies and fewer genes are fixed or present in a single genome (Figure 3II). In the limit where all genes are truly neutral, the mean frequency of all genes would be expected to be 0.5 (when the rates of gene gain and loss are equal). This is because each gene has equal chance of being present in any sampled genome, since we assume the population is at equilibrium and do not consider the effects of phylogeny and drift.

Many other possible DFEs among incoming genes can result in U-shaped gene frequency distributions arising in a population. Consequently, just changing the DFE results in a myriad of observed gene frequency distributions. For example, Figure 3III shows the simulated pangenome of 100 cells, each with 1000 genes whose fitness coefficients are sampled from an exponential distribution with rate parameter 1 and mean shifted to −1. In this case, most genes are highly deleterious, and many are not found in the simulated pangenome (Figure 3IIIa). In this case, most accessory genes are very rare, and only a small proportion of genes that are highly advantageous are present at high frequencies. These results illustrate that many distributions of fitness effects can potentially replicate the patterns observed in real pangenomes and would suggest that core genes are highly beneficial and accessory genes are mostly nearly neutral or slightly deleterious, in agreement with previous modelling results^16,28^. However, this conclusion also relies on simplifying assumptions that are unlikely to be met in the real world, such as equal rates of gene gain and loss for all genes.

#### Variation in gene gain and loss rates

Because the rates of gene gain and loss play an important role in determining the equilibrium frequency of a gene (as we showed in the previous section), we expect that their distributions will be important in shaping pangenomes. A reasonable approximation is that for most genes, the rates of gene gain and loss are very low, while for a small number of genes these rates can be relatively high. Thus, as an example, we sample these gain and loss rates from an exponential distribution. Figure 4 shows results from simulated pangenomes with fitness effects drawn from a normal distribution with mean 0 and standard deviation of 1 (blue density plot in Figures 4Ic, 4IIc and 4IIIc) and variable rates of gene gain and loss (Figures 4Ib, 4IIb and 4IIIb). If we fix the rate of gene loss, and draw the rates of gene gain from an exponential distribution with rate parameter 0.1 (Figure 4Ib), we find that some deleterious genes can be found at high frequencies (Figure 4Ie), when compared to the assumption of equal rates for all genes (Figure 3Id). If we fix the rate of gene gain and sample rates of gene loss from a similar exponential distribution (Figure 4IIb), we find an asymmetric U-shaped distribution in which many advantageous genes are now found at intermediate or low frequencies (Figure 4IIe). Finally, if we independently sample the three parameters for every gene (Figure 4III), we recover a symmetric U-shaped gene frequency distribution (Figure 4IIId). The U-shape of this distribution in this case is less extreme than that of Figure 3Ic, because there are more genes with intermediate frequencies for all values of fitness coefficient. This is due to both higher average gene gain and loss rates and the variation in these values. Overall, this analysis shows that if gene gain and loss rates vary for different genes, the observed gene frequencies are no longer determined by their fitness effects. Therefore, some beneficial genes may be present a low frequencies and deleterious genes at high frequencies.

**Figure 4.**
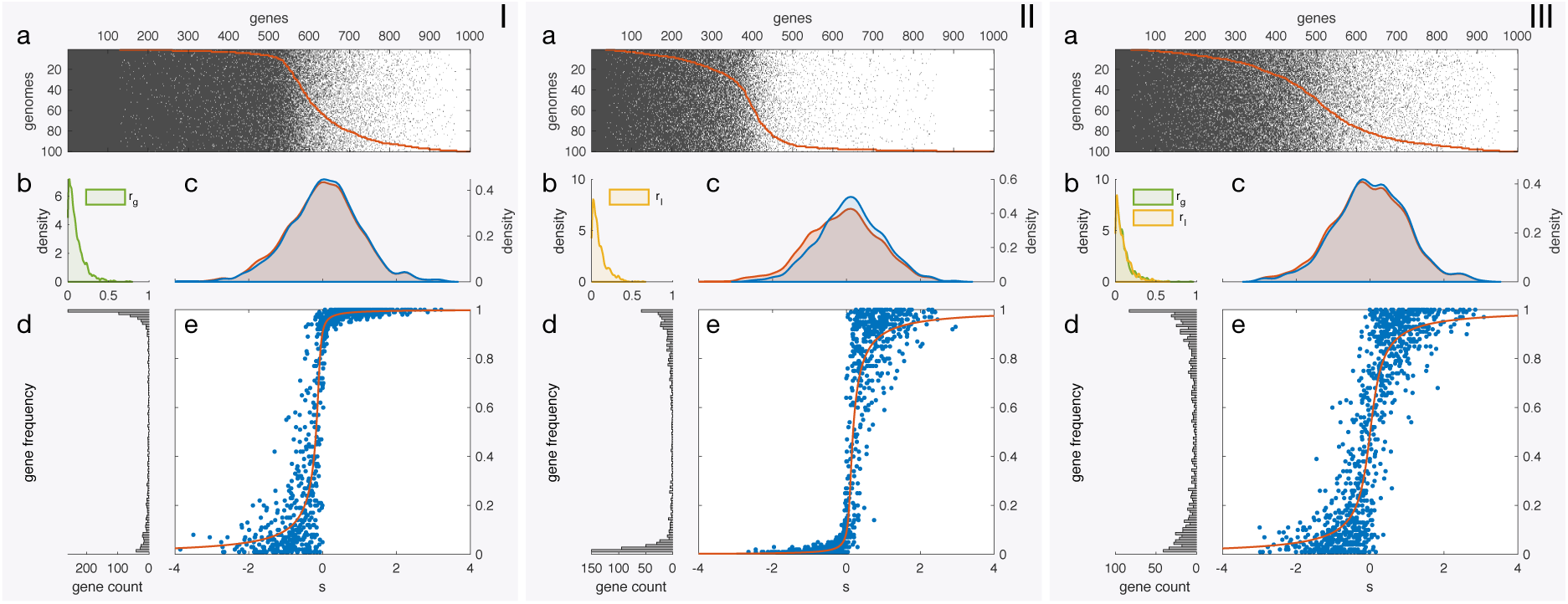
Simulated pangenomes for different distributions of rates of gene gain and loss. Each panel represents a simulation with rates of gene gain and loss sampled from the specified distribution. Fitness effects were sampled from a normal distribution with mean 0 and SD = 1. **I**. Rates of gain sampled from an exponential distribution with parameter 0.1 and fixed rate of loss (*r*_*l*_ = 0.01). **II**. Rates of loss sampled from an exponential distribution with parameter 0.1 and fixed rate of gain (*r*_*g*_ = 0.01). **III**. Rates of gain and loss sampled from independent exponential distributions with parameter 0.1. For each panel: **a**. Presence/absence of genes in the simulated pangenome, present genes are indicated in grey. The orange line indicates the number of strains in which a gene is present. **b**. Density plots for the sampled distributions of rates of gene gain and loss **c**. Density plots for the DFE of the gene pool (sampled from the specified theoretical distribution, in blue) and for the fitness effects of genes actually present in the pangenome (orange). **d**. Gene frequency distribution for the simulated pangenome. **e**. Gene frequencies versus fitness effects for 1000 genes in the simulated pangenome (blue dots) and the theoretical relationship (orange line).

### Niche-dependent fitness effects

The previous results show that if the focal organism can only live in a single ecological niche, then accessory genes are most likely to be either nearly neutral or deleterious, unless their rates of gene gain and loss are relatively high. However, besides the assumption of strong selection, this relies on the important but hidden assumption that a gene’s fitness effect is constant across all individuals of a given species. That is, the previous simulations have assumed that the contribution of a gene to a cell’s fitness is the same throughout the range of a species, in all possible environmental conditions and niches. This uniformity of fitness effects across environments is unlikely to be true for many genes^23^. For example, genes that can confer antibiotic resistance are likely to only be beneficial in environments where the antibiotic is present, and either confer no benefit or have a cost in its absence^29^. In other words, as the antibiotic concentration rises, presence of a resistance gene goes from normally deleterious (or neutral) to highly advantageous, to the point of potentially becoming essential under constant antibiotic pressure^30^.

To analyse the effect of niche dependence, we simulated pangenomes by sampling genomes from different ‘niches’, where a gene has different fitness effects, depending on the niche (Figure 5). In this case, to evaluate the *fitness contribution of a gene* to the simulated population sample, we introduce a new measure *ϕ*, which is the average fitness effect of the gene when present (see Methods). For example, if a gene is beneficial only in a particular niche and only present there, its fitness contribution will be positive. If it is also present in niches where its fitness effect is negative, its fitness contribution to the overall population will be reduced.

**Figure 5.**
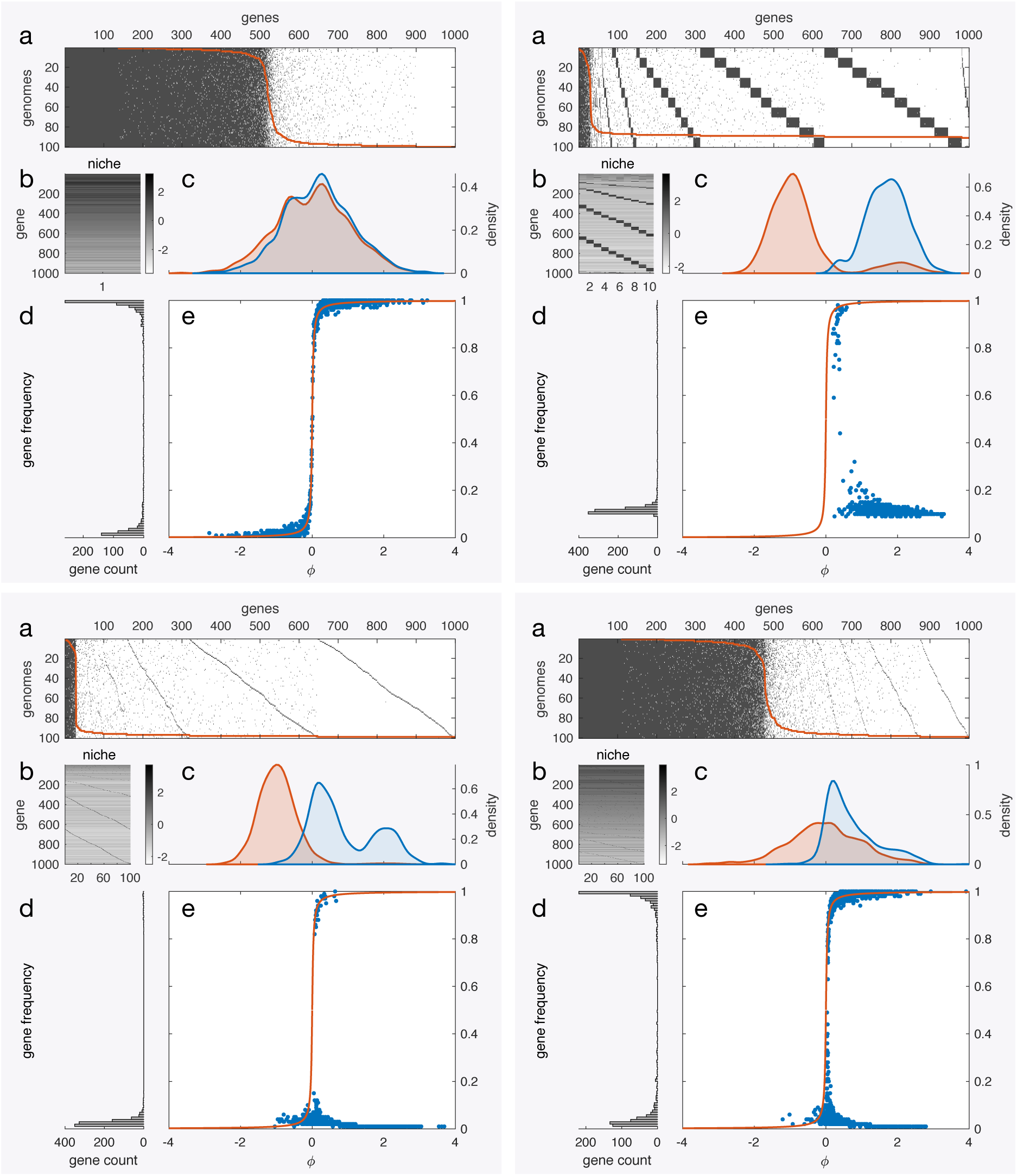
Simulated pangenomes for different distributions of rates of gene gain and loss. Each panel represents a simulation with a particular niche and population structure. **I**. 100 genomes from a single niche, 1000 genes with fitness effects drawn from a normal distribution with zero mean and unit standard deviation). **II**. 10 genomes simulated from each of 10 niches. 1000 mostly-deleterious genes (fitness effects drawn from a normal distribution with −1 mean and 0.5 standard deviation) can be gained in all niches. In each niche 100 genes (different for each niche) are mostly beneficial (fitness effect drawn form a normal distribution with mean 2 and 0.5 standard deviation). **III.** 1 genome simulated from each of 100 niches. 1000 mostly-deleterious genes (fitness effects drawn from a normal distribution with −1 mean and SD = 0.5) can be gained in all niches. In each niche 1 gene (different for each niche) is mostly beneficial (fitness effect drawn form a normal distribution with mean 2 and SD = 0.5). **IV**. 1 genome simulated from each of 100 niches. 1000 genes with fitness effects drawn from a normal distribution with zero mean and SD=1) can be gained in all niches. In each niche 1 gene (different for each niche) is mostly beneficial (fitness effect drawn form a normal distribution with mean = 2 and SD = 0.5). For each panel: **a**. Presence/absence of genes in the simulated pangenome, present genes are indicated in grey. The orange line indicates the number of strains in which a gene is present. **b**. Heatmap showing the sampled fitness effect of each gene in each niche **c**. Density plots for the DFE of the gene pool (sampled from the specified theoretical distribution, in blue) and for the fitness effects of genes actually present in the pangenome (orange). **d**. Gene frequency distribution for the simulates pangenome. **e**. Gene frequencies versus fitness effects for 1000 genes in the simulated pangenome (blue dots) and the theoretical relationship (orange line). Rates of gene gain and loss are assumed to be equal for all genes and niches (*r*_*g*_ = *r*_*l*_ = 0.01)

We first consider a single niche where gene fitness effects are drawn from a normal distribution as in the previous section. Figure 5Ia displays values of these fitness effects for 1000 genes as a heatmap. As before, this simulation recovers a U-shaped distribution of gene frequencies (Figure 5Id). In this case, the fitness contribution of a gene is identical to its fitness coefficient, because the fitness effect is the same for all genomes (Figure 5Ic). Again, we see that genes present at high frequency are on average highly advantageous (have a high fitness contribution) and rare genes or those present at intermediate frequencies are nearly neutral or deleterious. However, if we allow multiple niches, this situation can change drastically. Figure 5B shows an example where we consider 10 hypothetical niches. We assume that all genes in the gene pool are available for acquisition in all the niches, but genes are on average deleterious in 9 of the 10 niches and on average beneficial in just one. 100 genes are assumed to be on average beneficial in each niche, and we sample 10 genomes from each niche to simulate the pangenome shown in Figure 5IIa. In this case, most genes are only found in an average of 10 genomes (Figure 5IId, and square blocks in Figure 5IIa), those in which the genes are beneficial. Therefore, these relatively rare genes have a high fitness contribution (Figures 5IIe). In a reversal from the previous case, most genes found at high frequency make nearly neutral fitness contributions. Overall, all genes have a positive fitness contribution to the population (Figure 5IIc and 5IIe). By considering even more niches, for example 100 in Figure 5III and a single genome sampled from each, we obtain an even more extreme picture where all the genes that significantly contribute to fitness are very rare and most present in only one or two genomes (Figure 5IIId and 5IIIe). This result is in stark contrast to the simulations with constant fitness effects and low, uniform rates of gene gain and loss, where rare genes had a nearly neutral or deleterious effect. The situation corresponds to that described by McInerney et al.^12^, who proposed that where selection is highly effective (in species that have large long-term effective population sizes) and where multiple, diverse niches are occupied by the same species, we should expect accessory genes to accumulate and pangenomes to become quite large.

These simulations are extreme cases and intended as exemplars of the effect that niche-dependent selection can have on our expectations of the correlation between the frequency of a gene in the population and its fitness effect. Real pangenomes are more likely to be closer to the intermediate case presented in in Figure 5IV. In this case we also assume one genome sampled from each of 100 niches, where a single gene is likely highly beneficial, and the rest have fitness effects drawn from a normal distribution with mean zero. Therefore, the overall fitness effects of genes are similar to the single-niche situation presented in Figure 5A, except for the fact that each gene is likely to be highly beneficial in one niche. This simulation again recovers a U-shaped gene frequency distribution. The crucial difference in comparison to the single-niche case, is that now a substantial proportion of rare genes positively contribute to fitness in particular genomes (Figure 5IVc, 5IVd and 5IVe). This intermediate scenario then shows that accessory genes can be (nearly) neutral or advantageous.

In summary, our analysis shows that accessory genes present in pangenomes can be neutral, deleterious or advantageous, depending on particular properties of the genes, cells and environment. Therefore, the distributions and environment-dependence of gene fitness effects and rates of gene gain and loss are thus crucial for understanding the composition of pangenomes.

## Discussion

We have presented a model to simulate prokaryote pangenomes that allows us to analyse the effect of important parameters on gene frequencies and fitness contributions. We have shown that fitness effects, rates of gene gain and loss and their respective distributions are crucial in determining pangenome properties. Depending on assumptions and circumstances, the model shows that accessory genes can be neutral, advantageous or even deleterious. Therefore, this model contributes to reconcile previous models which suggest that pangenomes are the result of purely neutral or purely selective processes, and that accessory genes are mostly neutral or beneficial^11,15–19^. Our aim is to highlight factors that may be important in shaping pangenomes and that perhaps have received little attention, and therefore our model is intended as a framework to investigate how basic parameters can affect gene content, rather than a detailed account of pangenome evolution. By assuming a regime with strong selection and disregarding drift, our model lies at the opposite end to models that assume pure neutrality in the evolutionary process. This assumption may be justified for the large population sizes of many prokaryotes in which selection is expected to be very effective and drift may be relatively weak, as suggested by high levels of codon adaptation in some species and the general compactness of prokaryote genomes^12^. Together with the assumption of equilibrium for gene frequencies, these simplifications allow a flexible framework where we can incorporate many of the processes that influence pangenome properties.

It is known that the distribution of fitness effects of mutations or genes is a major determinant of the levels of genetic polymorphism we should expect in populations^8,27^, and we have shown how this can impact the gene frequency distribution (Figure 3). Disregarding population structure and variation in the rates of gene gain and loss, we find that as the proportion of neutral and deleterious genes in the DFE increases, the proportion of low frequency genes in the pangenome also increases. However, we show that the DFE is not the only determinant of gene frequencies, and that rates of gene gain and loss can have a large influence on gene content and on the types of genes we expect to find in pangenomes^31^. If these rates are low relative to fitness effects, selection can maintain advantageous genes essentially fixed in the population, and genes present at low frequencies will be nearly neutral. On the other hand, if these rates are high, deleterious genes can be common in the population, while advantageous genes may not reach fixation. Although, in reality, these rates may on average be low with respect to fitness effects, they are likely to vary for different genes, genomes and species, and the values of these parameters and their distributions may contribute significantly to gene content variation in prokaryote genomes. In particular, genes with high gain rates can be maintained in the population at high frequencies even if they are significantly deleterious. This is likely the case for some MGEs and lysogenic viruses, what we call ‘infectious’ genes, which are most likely deleterious but gained at high rates. For example, deleterious polymorphisms can become prevalent in a bacterial population due to the effect of high recombination mediated by conjugative plasmids^32^, and inteins with high fitness costs seem to be maintained in genomes^33^.

Furthermore, the genetically-determined component of the rates of gene gain and loss could itself be a target of natural selection, from the perspective of either the receiving cell or an infectious gene. Besides selection on the rates of natural transformation and DNA recombination, mechanisms that quickly recognise and destroy foreign DNA, such as restriction modification and CRISPR systems may not only prevent viral infections, but also reduce the rate of gene gain in cells that possess these systems, regardless of their fitness effects. ‘Addictive genes’ are another example, where a gene (or operon), even if deleterious, once acquired becomes essential and is difficult to lose. Such may be the case of anti-toxin genes of toxin/anti-toxin systems and restriction enzymes in restriction modification systems^34,35^. Another possible example of beneficial change in the loss rate of a gene may be the expansion of transposon numbers within a genome, and the observed loss of sequences responsible for transposition for a widely-spread gene encoding colistin resistance^36^.

We have also highlighted the potential of niche-dependent fitness effects for maintaining variability in gene content. Although we lack information of how prevalent these effects are, we know that individuals from the same prokaryote species can survive and thrive in very different environments^37^. Therefore, it is likely that these fitness by environment interactions for particular genes are relatively common. Besides previously mentioned antibiotic genes, certain metabolism genes are good candidates for having niche-specific benefits^38^. Examples are the association of vitamin B5 synthesis genes with cattle-specific strains of *Campylobacter*^39^, and the acquisition by HGT of transporter-encoding genes with environment-dependent fitness effects in yeasts^40^. Moreover, we do not really know how environments are partitioned, and what may appear as a homogeneous niche may in fact have distinct compartments with specific selective pressures. For example, pathogens multiplying in different organisms, parts of the body and even different parts of the same organ may be subject to different conditions in terms of nutrients, physicochemical properties and immune responses^41,42^. Even apparently homogeneous environments such as the open ocean are more heterogeneous than previously thought, having niches which vary significantly in availability of resources and other environmental conditions^43^. In any case, the existence of these ‘gene by environment’ effects could account for the presence of accessory genes that may be highly advantageous under some, but not all, conditions. In fact, even gene essentiality can be environment-dependent^44^.

Overall, we argue that for an individual gene, its particular circumstances and details matter, and therefore its contribution to fitness cannot be extrapolated from its population frequency alone. However, in general, neutral or deleterious genes are unlikely to be part of the core genome, unless populations are subjected to significant genetic drift, their gain rates are very large or/and their loss rates very small. Besides these broad conclusions, it is currently difficult to make strong claims as to whether accessory genes are on average advantageous or not. Our model shows that very different assumptions can broadly replicate the patterns of gene presence/absence we observe in nature. Therefore, the issue is not that the data do not fit our models, but that we lack information to truly distinguish between the different possible explanations. If we want to understand the average properties of pangenomes, we need more information about the distributions of fitness effects and rates of gene gain and loss. Furthermore, it is worth noting that even though we have simulated parameter values as independent, this will not necessarily be the case for real genes. For example, genes that are highly beneficial are also likely to be more abundant in the environment and may have higher gain rates. On the other hand, deleterious genes present in MGEs might also be gained at high rates. Since we lack information about the individual distributions of these parameters, it is not surprising that we know little about their joint distributions.

In conclusion, further understanding of the existence and maintenance of pangenomes in different prokaryote groups, requires not only genome sequences, but extensive metadata, environmental parameters and phenotypic characteristics. We need to understand the extent and nature of the structure of bacterial populations, and how much gene flow there is between subpopulations. Finally, we need longitudinal information about how gene frequencies change in the population, and the stability of particular polymorphisms and pangenomes in general. Collecting all this information not only requires extensive environmental sampling but will also likely rely on experiment inside and outside the lab to determine the values of parameters under controlled conditions.

## Methods

### Calculation of gene frequencies

From Eq. 1 in the model description, we obtain the steady state gene frequency, *x*_*ss*_, which is described by a quadratic equation with one positive and thus meaningful solution:

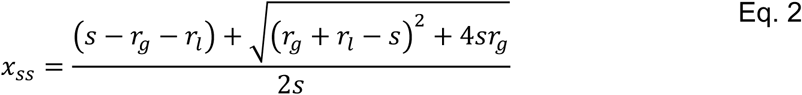

We use Equation 2 to calculate the expected frequency of a gene in the population. Although *x*_*ss*_ is undefined when *s* = 0, this equation can be used to calculate equilibrium frequencies for nearly neutral genes, with fitness effects arbitrarily close to zero. The equilibrium assumption may not be entirely justified in real populations (in particular if gene gain and loss rates are low and fitness effects are low), but it is a useful assumption that allows us to explore how parameters affect gene frequencies and their distributions for large numbers of genes. An analytic solution describing the temporal dynamics of gene frequency also exists for Equation 1 (see supplementary information).

### Pangenome simulations

To simulate pangenomes, we assume there is a finite ‘gene pool’, a set of *n* genes that can be gained and lost from cells in the population (Figure 1b). If we make the simplifying assumption that there is no interaction between genes, the gene frequency model described above can be used to model multiple independent genes in the population and thus explore the effects of model parameters on pangenomes. This assumption implies that there are no epistatic effects between genes, and thus the parameters describing the frequency of one gene are independent from those describing any other gene (Also see Supplementary Information for derivation of the same equation from genotypes for two genes). For each simulation, we assume values for each of the three model parameters (*s, r*_*g*_, *r*_*l*_) for each gene. Depending on the simulation, the values can be identical for all the genes, or sampled from a specified distribution. Using these values, we calculate the expected equilibrium frequency for each gene using Equation 2. In order to simulate the presence of each gene in each of *N* genomes, a uniformly-distributed random number in the [0,1] interval is generated, and the presence of the gene is accepted if the random number is smaller than the calculated equilibrium gene frequency.

### Simulations with niche-dependent fitness effects

To simulate pangenomes from different niches, we first define a general gene pool with *n* genes whose fitness coefficients are drawn from a specified distribution (usually with negative average values). In each of *k* niches the fitness effects of the genes are assumed to be the same as the generalised gene pool, except for a defined subset of genes whose fitness effects are drawn from a different distribution (usually with positive average values). In the simulations presented in Figure 5 we consider non-overlapping sets of *n/k* genes to have niche-dependent fitness values. Finally, we simulate a specified number of genomes from each niche as described above.

### Fitness contribution of a gene

To define the average contribution of a gene to the population fitness, *ϕ*, we consider a sample of N genomes. For a particular gene, we define *z*_*i*_ as a variable representing the presence or absence of a gene in genome *i*, where its value is equal to 1 if the gene is present, and 0 otherwise. In this case, *s*_*i*_ represents the fitness coefficient of the gene in the niche from which genome *i* is sampled. The fitness contribution of a gene *ϕ* is then defined as:

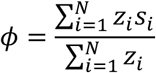

## Acknowledgements

We thank John Brookfield, Caroline Colijn, Jessica Stockdale and Fiona Whelan for comments on the manuscript and members of the O’Connell and McInerney groups for discussion. JOMc is supported by a BBSRC Responsive Mode Grant BB/N018044/2.

