## Supplementary for "Selection-based model of prokaryote pangenomes"

Analytic solution

The differential equation for our model of gene frequency:

$$\frac{dx}{dt} = r_g(1-x) + sx(1-x) - r_l x = r_g + (s - r_g - r_l)x - sx^2$$

as the following analytical solution:

$$x(t) = \frac{\sqrt{4ac-b^2}}{2a}\tan\left[\frac{t\sqrt{4ac-b^2}}{2a} + \tan^{-1}\left(\frac{2ax_0+b}{\sqrt{4ac-b^2}}\right)\right] - \frac{b}{2a}$$

where $x_0$ is the gene frequency at $t = 0$, and

$$a = -s$$
$$b = s - r_g - r_l$$
$$c = r_g$$

Derivation of model from two-gene genotypes

Assume population with two accessory genes $a$ and $b$, with fitness coefficient $s_a$and $s_b$, respectively. These genes can be gained or lost from the genome with rates $r_g$ and $r_l$ respectively. 4 genotypes in the population, each at frequency $y_{ba}$, such that:

$$y_{00} + y_{01} + y_{10} + y_{11} = 1$$

The allowed transitions between genotypes are described by the diagram below:

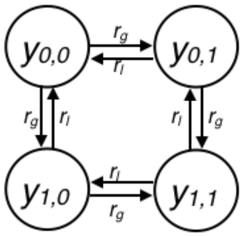


Therefore, the rates of change of each genotype are described by a differential equation:

$$\frac{dy_{00}}{dt} = r_l(y_{01} + y_{10}) - 2r_g y_{00} + \mu y_{00} - k_T y_{00}$$
$$\frac{dy_{01}}{dt} = r_g y_{00} + r_l y_{11} - (r_g + r_l)y_{01} + (\mu + s_a)y_{01} - k_T y_{01}$$
$$\frac{dy_{10}}{dt} = r_g y_{00} + r_l y_{11} - (r_g + r_l)y_{10} + (\mu + s_b)y_{10} - k_T y_{10}$$
$$\frac{dy_{11}}{dt} = r_g(y_{01} + y_{10}) - 2r_l y_{11} + (\mu + s_a + s_b)y_{11} - k_T y_{11}$$

where $\mu$ is the population growth rate, and

$$k_T = \mu y_{00} + (\mu + s_a)y_{01} + (\mu + s_b)y_{10} + (\mu + s_a + s_a)y_{11}$$

So that the population size is constant.

Summing over the genotypes to obtain gene frequencies $x_a$ and $x_b$, that is: $x_a = y_{01} + y_{11}$, and $x_b = y_{10} + y_{11}$, and assuming there is no interaction between genes (so $y_{11} = x_a x_b$), it is possible to obtain a differential equation for the frequency of each gene $i$, where $i$ denotes $a$ or $b$:

$$\frac{dx_i}{dt} = r_g(1 - x_i) + s_i x_i(1 - x_i) - r_l x_i$$

Therefore, the frequency of each gene is independent from the other (and independent of μ
A similar result is valid for more than two genes, and we can use this differential equation to analyse the frequency of many genes independently

### Supplementary Figures

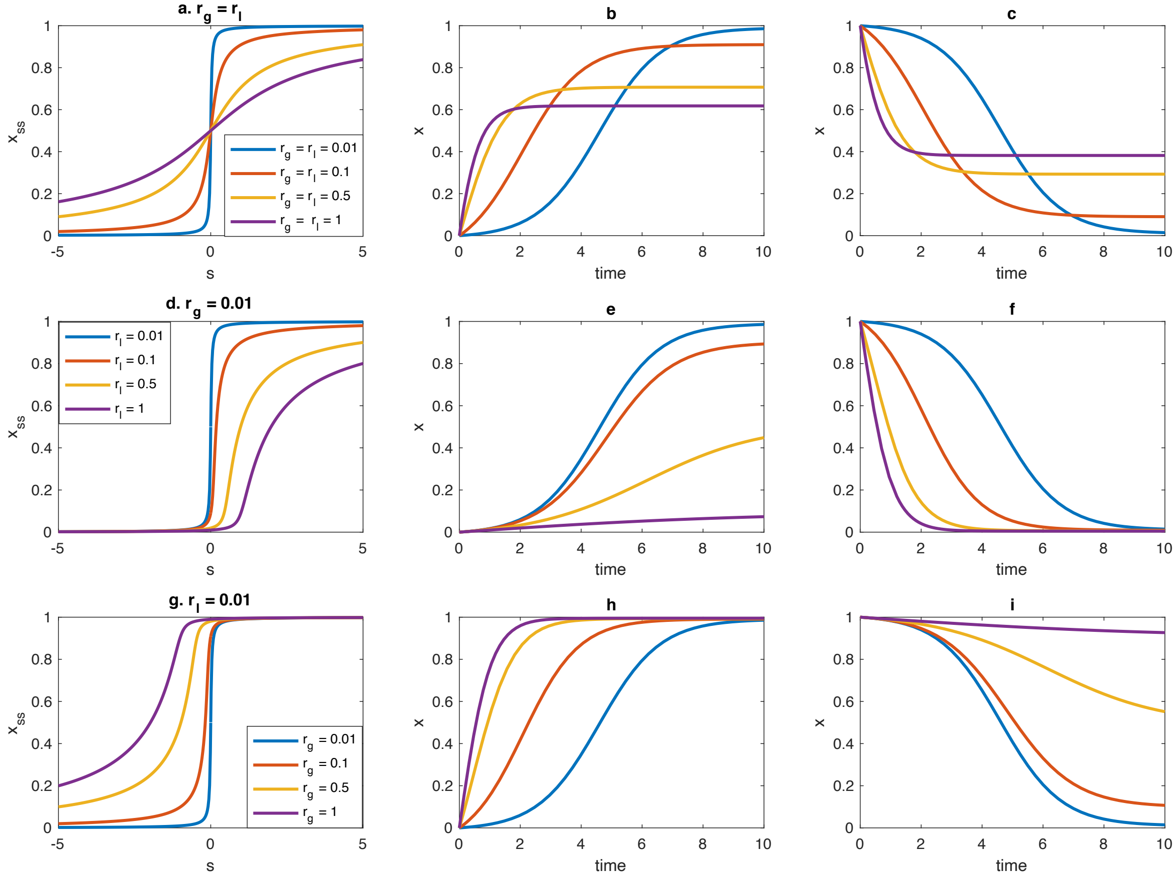


***Supplementary Figure 1***. Dependence of gene frequency and dynamics with respect to model parameters. Left column shows the dependence of gene frequency with respect to the fitness effect, $s$, for different rates of gene gain and loss. The middle column shows the dynamics of gene frequency for a gene with $s = 1$ starting at $x = 0$. The right column shows the dynamics of gene frequency for a gene with $s = -1$ starting at $x = 1$. First row (a, b, c), effect of rates of gene and loss when $r_g = r_l$. Second row (d, e, f), effect of the rate of gene loss with fixed rate of gene gain ($r_g = 0.01$). Third row (g, h, i), effect of the rate of gene gain with fixed rate of gene loss ($r_l = 0.01$).

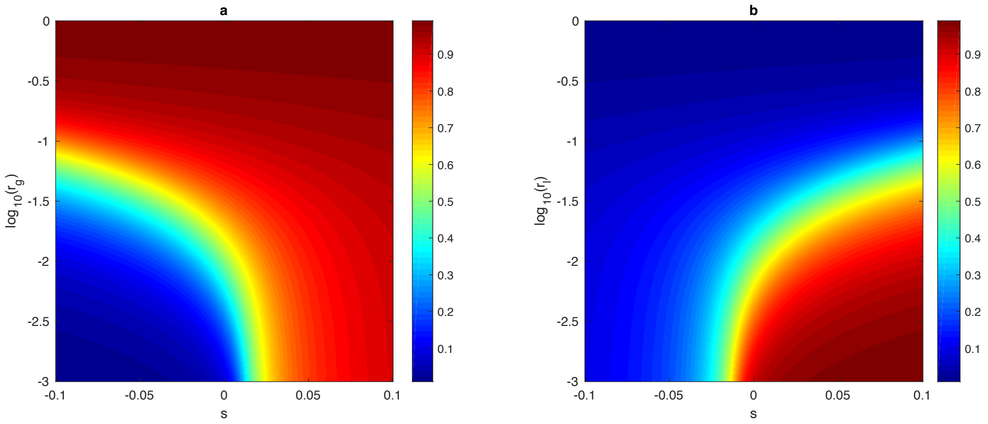


***Supplementary Figure 2***. Gene frequencies with respect to rates of gene gain (a) and loss (b) for different values of fitness effects. Colour bar indicates the equilibrium gene frequency $(x)$. For (a) $r_l = 0.01$, and for (b) $r_g = 0.01$.
